# Non-intravenous, carbapenem-sparing antibiotics for the treatment of bacteremia due to ESBL or Amp-C β-lactamase: A propensity score study

**DOI:** 10.1101/304246

**Authors:** Y. Meije, C. Pigrau, N. Fernández-Hidalgo, M. Clemente, L. Ortega, X. Sanz, J. Loureiro-Amigo, M. Sierra, A. Ayestaran, A. Morales-Cartagena, A. Ribera, J. Rodríguez-Baño, J. Martínez-Montauti

**Affiliations:** infectious Diseases Unit – Internal Medicine Department. Hospital de Barcelona. Societat Cooperativa d’Installacions Assistencials Sanitàries (SCIAS). Barcelona, Spain.; Servei de Malalties Infeccioses. Hospital Universitari Vall d’Hebron. Universitat Autònoma de Barcelona, Barcelona, Spain.; Spanish Network for the Research in Infectious Diseases (REIPI RD12/0015), Instituto de Salud Carlos III, Madrid, Spain.; Microbiology Department. Hospital de Barcelona. Societat Cooperativa d’Installacions Assistencials Sanitàries (SCIAS). Barcelona, Spain.; Pharmacy Department. Hospital de Barcelona. Societat Cooperativa d’Instal·lacions Assistencials Sanitàries (SCIAS). Barcelona, Spain.; Unidad Clínica de Enfermedades Infecciosas, Microbiología y Medicina Preventiva. Hospital Universitario Virgen Macarena / Departamento de Medicina, Universidad de Sevilla / Instituto de Biomedicina de Sevilla, Seville, Spain.

**Author notes:** Corresponding author, Yolanda Meije, MD, PhD., Infectious Disease Unit – Internal Medicine Department, Hospital de Barcelona. SCIAS Diagonal 660, 08034, Barcelona.

**Keywords:** Carbapenem-sparing antibiotics, Enterobacteriaceae bacteremia, stewardship, extended-spectrum β-lactamase, Amp-C β-lactamase, cotrimoxazole, trimethoprim-sulfamethoxazole, quinolones, ciprofloxacin.

## Abstract

**Introduction:** Carbapenems are considered the treatment of choice for extended-spectrum β-lactamase (ESBL) or Amp-C β-lactamase-producing Enterobacteriaceae bacteremia. Data on the effectiveness of non-intravenous carbapenem-sparing antibiotic options are limited.

**Objective:** To compare the 30 day-mortality and clinical failures associated with the use of carbapenems vs an alternative non-intravenous antibiotic for the definitive treatment of ESBL/Amp-C positive Enterobacteriaceae bacteremia.

**Methods:** This is a 12-year retrospective study (2004 – 2015) including all patients with bacteremia due to ESBL/Amp-C-producing Enterobacteriaceae. Given the lack of randomization of the initial therapies, a propensity score for receiving carbapenems was calculated.

**Results:** There were 1115 patients with a first episode of bacteremia due to *E. coli* or *K. pneumoniae*, of which 123 were ESBL/Amp C-positive (11%). There were 101 eligible patients: 59 in the carbapenem group and 42 in the alternative treatment group (cotrimoxazole 59.5%, quinolones 21.4%). The most frequent sources of infection were urinary (63%) and biliary (15%). Compared to the carbapenem group, patients treated with the alternative regimen had a shorter hospital stay (median [IQR]: 7 days [5-10] vs 12 days [9-18], p<0,001). The use of an alternative non-IV treatment did not increase mortality (OR 27; 95% CI 0.05-1.61; p=.15). After controlling for confounding factors with the propensity score, the adjusted OR of carbapenem treatment was 4.95; 95% CI (0.9426.01, p=.059).

**Conclusion:** Alternative non-IV carbapenem-sparing antibiotics could have a role in the definitive treatment of ESBL/Amp-C-positive Enterobacteriaceae bloodstream infections, allowing a reduction in carbapenem use. The use of cotrimoxazole in this setting has shown favourable results.

ESBL: Extended-spectrum β-lactamase
Amp-C: Amp-C β-lactamase
BL/BLIs: β-lactam/β-lactamase inhibitor combinations
IV: Intravenous
TMP-SMX: Trimethoprim-Sulfamethoxazole

Some of the data contained in this article were presented at the 55^th^ Interscience Conference on Antimicrobial Agents and Chemotherapy (ICAAC) and at the 28^th^ International Congress of Chemotherapy Meeting (ICC), San Diego, USA, 2015.

## INTRODUCTION

Extended-spectrum β-lactamase (ESBL) or Amp-C β-lactamase (Amp-C)-producing Enterobacteriaceae have been increasingly implicated in health care– and community-associated bacteremia (1). Effective treatment of ESBL or Amp-C bacteremia has become a major challenge due to frequent resistance to various antibiotics and the existence of mechanisms of co-resistance in this setting (2). At present, carbapenems are the treatment of choice for ESBL bacteremia (3). However, increasing carbapenem resistance among Enterobacteriaceae, as well as in other bacteria, calls for a more judicious approach to carbapenem use (4).

Previous studies of empiric treatment of ESBL Enterobacteriaceae bacteremia with beta-lactam/beta-lactam inhibitor combinations (BL/BLIs) are contradictory (3, 5-7). These discrepant results may be due to differences in the source of infection, the genetic background of the microorganism, or local epidemiology (8). Nevertheless, evidence is emerging that empiric or definitive treatment with BL/BLIs is probably as effective as carbapenem therapy in the setting of ESBL *(E. coli* or *K. pneumoniae)* bacteremia (9-11). Data regarding the usefulness of carbapenem-sparing antibiotics other than BL/BLIs for definitive treatment have also been reported, though also within an IV regimen (12-14). The effectiveness of non-intravenous (oral or intramuscular) antibiotic treatment for the management of ESBL or Amp-C bacteremia has not been widely assessed to date (3, 15). At our cooperative non-profit private hospital, patients with ESBL or Amp-C Enterobacteriaceae bacteremia frequently undergo definitive treatment with an orally administered antimicrobial agent. This hospital encourages close follow-up of patients who may re-contact their doctor directly if fever or other signs of possible infection appear after discharge.

The aim of this study was to compare the 30-day mortality and clinical failure in two groups: patients receiving carbapenems *vs* an alternative therapy, based on a non-IV carbapenem-sparing antibiotic regimen, for the definitive treatment of ESBL or Amp-C-ositive Enterobacteriaceae bacteremia.

## METHODS

### Study design, setting and participants

This 12-year retrospective study (January 2004 – December 2015) was conducted at a tertiary general hospital, with 250 beds in Barcelona, Spain. Patients over age 15 with community-acquired or healthcare-associated bacteremia due to ESBL or Amp-C-producing Enterobacteriaceae were included. Patients who died in the first 72h or those without one-month follow-up were excluded. If patients experienced more than one bacteremic episode, only the first episode was included. We recorded the prescribed antibiotic in each case, as selected by the patient’s attending physician. All episodes were identified from the electronic microbiological database. The patients’ clinical information was collected from electronic clinical charts and electronic pharmacological database. The follow up was performed by either the electronic clinical charts or by telephone if the patient had been discharged. This study was approved by the institutional review board for clinical trials.

### End-points

The primary outcome measure was the 30-day mortality rate, and the secondary outcomes were clinical failure within 30 days of onset of bacteremia and length of hospital stay.

### Variables and Definitions

Bacteremia was defined as the isolation of organisms in one or more separately obtained blood culture with compatible clinical features. The cases of bacteremia were categorized as nosocomial, healthcare-associated, or community-acquired in accordance with the criteria of Friedman et al^16^. Infections were defined as urinary tract, biliary, incisional wound, soft-tissue, catheter-related or primary bloodstream infection, in accordance with the Centres for Disease Control and Prevention guidelines (16). The following patients were considered immunocompromised: those receiving corticosteroids at a dose of ≥20 mg prednisone or equivalent for ≥2 weeks, those with neutropenia (absolute neutrophil count below 500/mm3) or those receiving anticancer chemotherapy in the previous six months. Chronic kidney disease (CKD) was defined and staged according to the Kidney Disease Improving Global Outcomes definition and classification (17). Charlson comorbidity score was defined as previously described by Charlson et al (18). The severity of bacteremia on the day of onset was graded with the Pitt bacteremia score (19). Source control was defined as any kind of intervention apart from antibiotic treatment applied to solve the infection, such as surgical treatment, abscess drainage or catheter withdrawal. Sepsis or septic shock was defined according to current definitions (20).

Antimicrobial therapy was regarded as empirical if administered before the susceptibility test results were available. Modification of treatment was defined as a change to an active antibiotic after the culture result became available, in accordance with the pathogen’s *in vitro* susceptibility pattern. Definitive therapy was defined as an active antibiotic administered for > 50% of the total duration of antimicrobial therapy after the antibiogram result. Treatment was defined as appropriate when an active antimicrobial agent, determined by *in vitro* susceptibility testing, was administered at the usual recommended dose. Clinical failure was defined as persistence of bacteremia (i.e. positive blood cultures for the same Enterobacteriaceae after 72 hours of active antibiotic treatment by *in vitro* susceptibility), persistence of fever or sepsis, death, or relapse during a 30-day follow-up, defined as positive blood cultures for the same microorganism (after a previous negative result). Length of stay was defined as the time from the first positive blood culture to discharge.

### Microbiological analysis

Microbiological identification and antibiotic susceptibility tests were carried out using the MicroScan WalkAway system (Beckman Coulter, Inc., Brea, CA). Presence of ESBL and/or AMPc was screened in all isolates with diminished susceptibility to cephalosporins by Microscan System, and confirmed by double disc synergy test (DDST), combination disc test, gradient test method, or molecular characterization by PCR, according to CLSI (Clinical & Laboratory Standards Institute) and EUCAST (European Committee on Antimicrobial Susceptibility Testing) guidelines. The β-lactams used for confirmation, testing their synergistic effect with amoxicillin-clavulanate were: ceftazidime, cefotaxime, and aztreonam. The following β-lactams were used for confirmation, testing their synergistic effect with amoxicillin-clavulanate: ceftazidime, cefotaxim and aztreonam.

During 2004-2006 period, in vitro susceptibility tests were interpreted based on the CLSI breakpoints (Clinical and Laboratory Standards Institute: M100: Performance Standards for Antimicrobial Susceptibility Testing) (21), and during 2007-2015 on the EUCAST breakpoints (European Committee on Antimicrobial Susceptibility Testing) (22).

During the study period, the microbiology department at our hospital did not have access to PCR for the study of Amp C beta-lactamase-producing *Escherichia coli*. As we were unable to establish whether *E.coli-AmpC* enzymes were encoded by chromosomal or plasmid genes, we excluded all AmpC-*E. coli* to prevent potential confounding.

### Statistical analysis

Quantitative variables were expressed as medians and interquartile ranges (IQR); categorical variables were reported as absolute numbers and percentages. To detect significant differences between groups, we used the Chi-square test or Fisher exact test for categorical variables, and the Student t test or Mann-Whitney U test for continuous variables, as appropriate. Independent predictors for 30-day mortality were identified by logistic regression analysis.

Given the lack of randomization of the initial therapies, a propensity score for receiving carbapenems was estimated using a backward stepwise logistic regression model that included variables with P values ≤25 in the univariate analysis, plus other variables considered relevant in deciding the empiric treatment. The following variables were included: age, sex, Pitt index, active cancer, chronic kidney disease, source of bloodstream infection, empiric treatment (as appropriate or not) and the time without effective treatment. An Inverse probability of treatment weighting (IPTW) logistic regression using the propensity score was fitted to estimate the risk of mortality due to carbapenem administration. The weights to the propensity score were finally obtained after fitting a logistic regression model for use of carbapenem as outcome. The model obtained had an Area under ROC curve of 0.77.

The statistical analysis was conducted using SPSS software for Windows, version 17.0 (SPSS Inc., Chicago, IL, USA) and STATA 13.1 (Statacorp College Station, Tx, USA)

## RESULTS

During the study period, there were 1309 patients with a first episode of bacteremia due to Enterobacteriaceae (1115 due to *E. coli* or *K. pneumoniae)* of which 123 (11%) were ESBL or Amp C-positive *E. coli* or *K. pneumoniae*. Twenty two patients were excluded (Figure 1), resulting in a final cohort of one hundred and one patients, which were grouped as per type of treatment: 59 in the carbapenem group and 42 in the alternative non-IV treatment group (TMP-SMX 25, quinolones 9, aminoglycosides 5, fosfomycin 2, amoxicillin/clavulanate 1).

**Figure 1.**
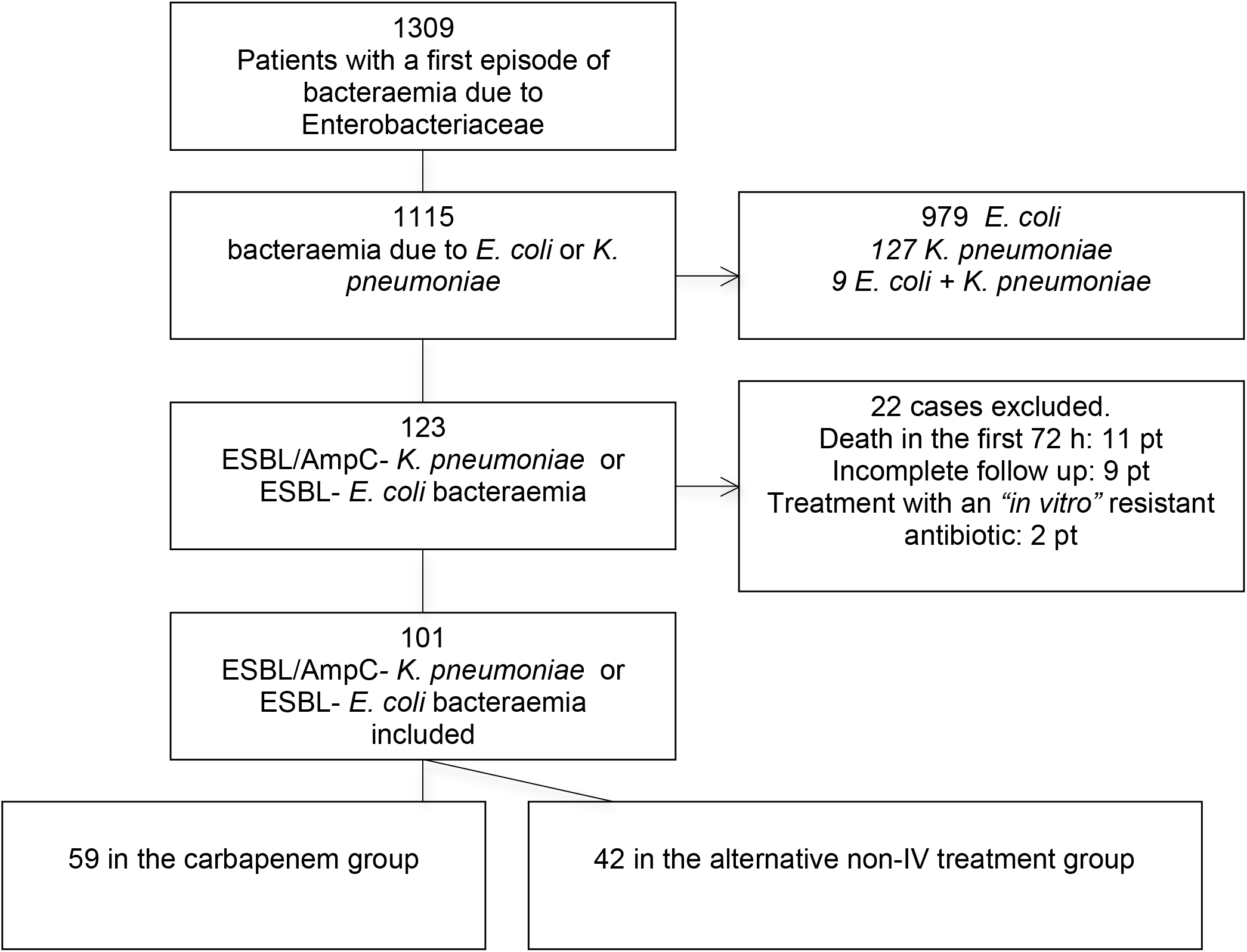
Flow chart.

The in vitro susceptibility rate for various antibiotics for ESBL/Amp-C-producer strains was as follows: carbapenem 100%, aminoglycosides 76%, piperacillin/tazobactam 59%, TMP-SMX 38%, amoxicillin/clavulanate 27% and quinolones 14%.

The most frequent infection sources were urinary (63%), biliary (15%) and unknown source (8%), followed by catheter-related (6%), intra-abdominal (5%), surgical wound/soft tissues (2%) and prosthetic joint infection (1%). The clinical and demographic characteristics of each group are shown in Table 1. There were no differences between groups (carbapenems *vs* alternative therapy) in terms of age, comorbidity, infection source, severity of underlying disease, time of empiric or definitive treatment. Compared to the carbapenem group, the patients treated with the alternative regimen had lower median Pitt score.

**Table 1.**
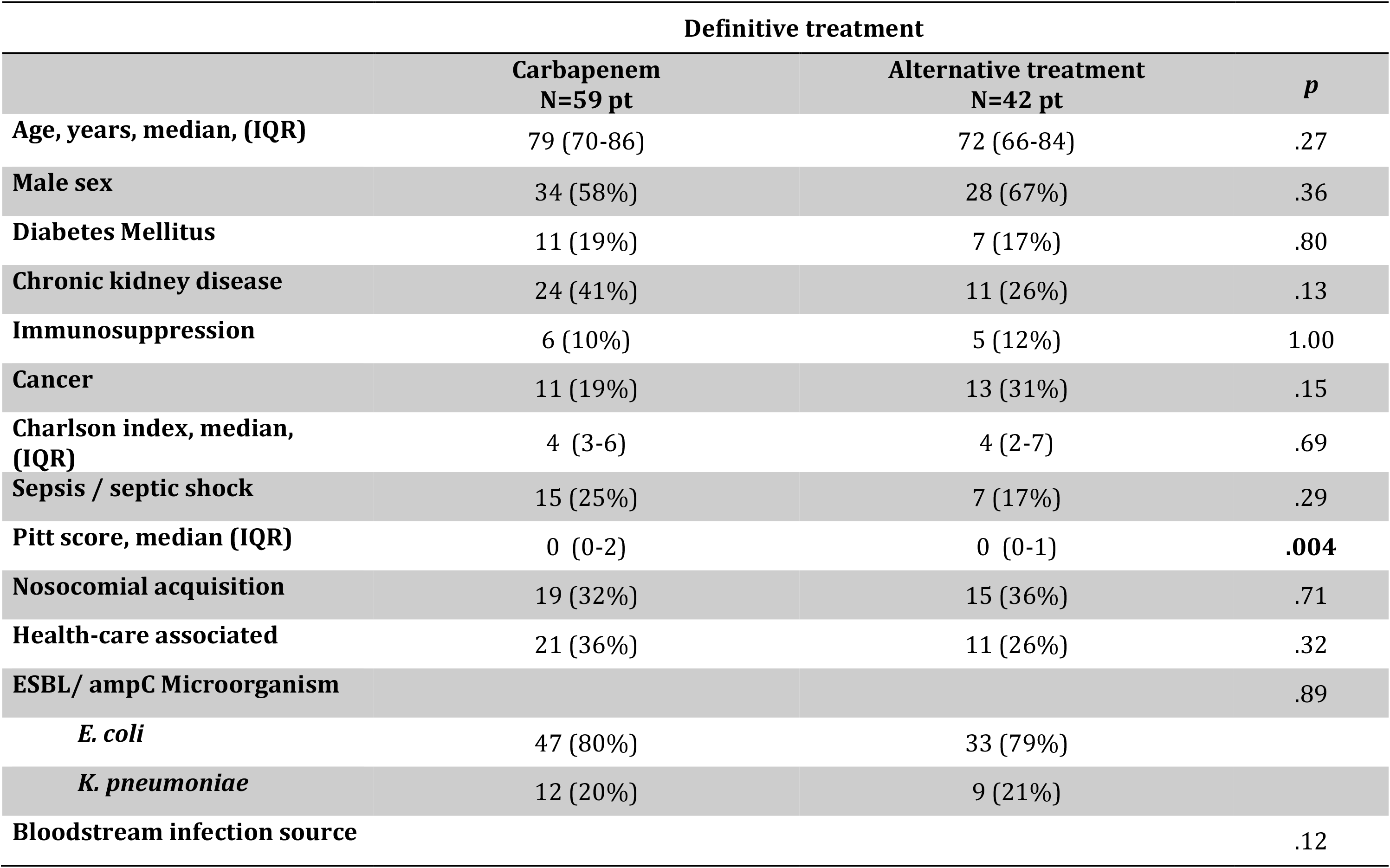
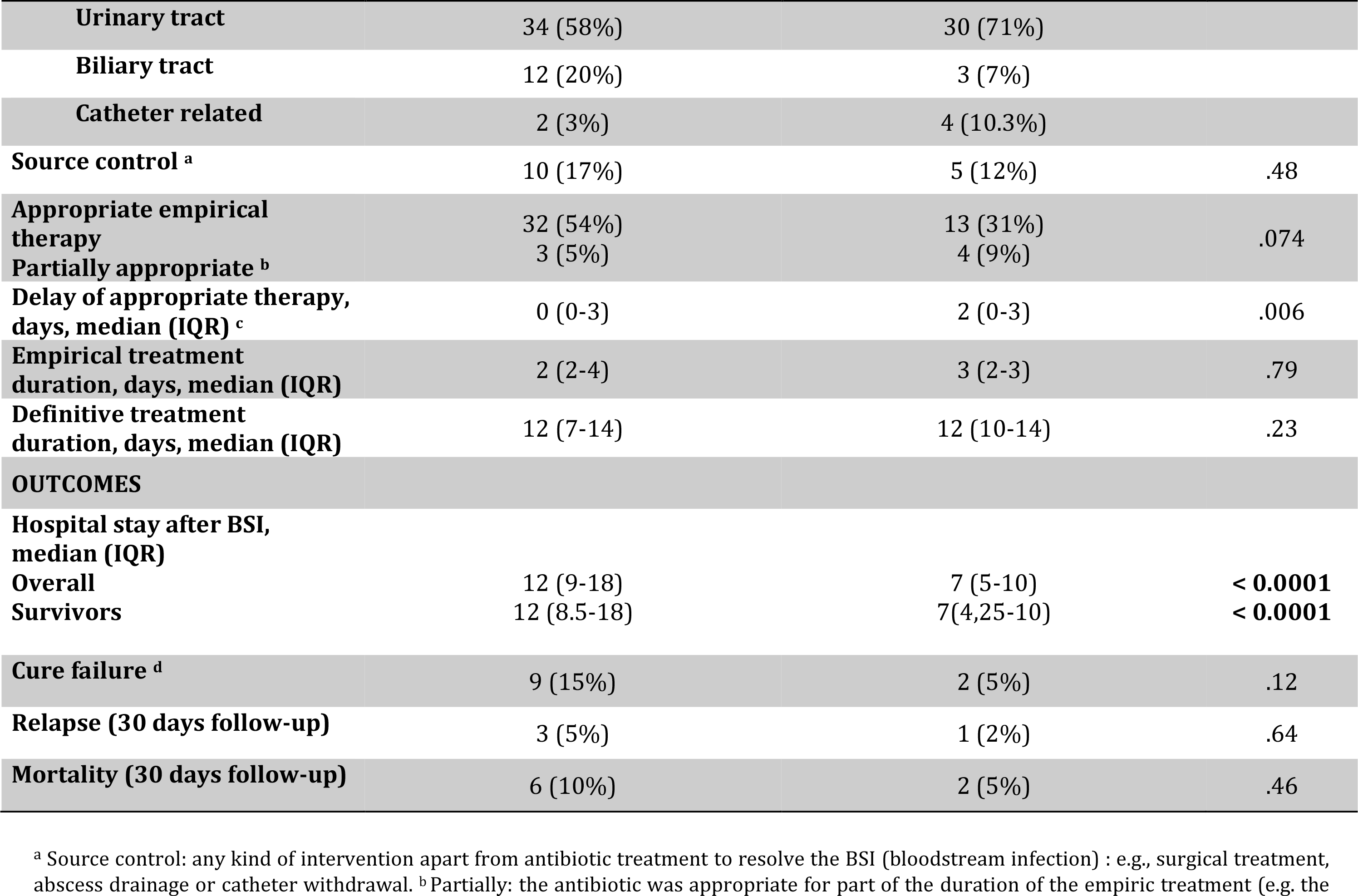
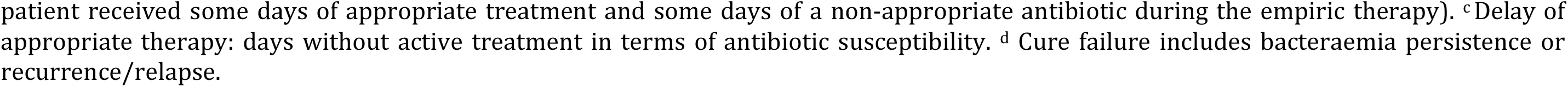
Characteristics of patients with bloodstream infections caused by extended-spectrum b-lactamase-producing Enterobacteriaceae according to definitive therapy.

Source control was performed in five patients out of 42 (12%), in the alternative group: three underwent catheter removal due to a catheter-related bacteremia and two underwent endoscopic retrograde cholangiopancreatography due to a bacteremia of biliary source. Source control was performed in ten patients out of 59 (17%), in the carbapenem group: three underwent endoscopic retrograde cholangiopancreatography, five required double J catheter or percutaneous nephrostomy, one required debridement and implant retention, and one needed an abdominal surgery.

During the 30-days of follow up, among the 59 carbapenem treated patients, 6 (10%) died and 9 (15%) were considered as clinical failure (including the 6 patients who died), of which 3 were due to bacteremia relapse. In the non-carbapenem group, two patients (5%) died, which were also considered as clinical failure, and one of them had also previously developed bacteremia relapse (Table1). The two patients who died in the alternative group had a disseminated cancer (bladder and colon cancer). Among the 6 patients who died in the carbapenem group one had a pancreas tumor and the rest of them were patients with multiple comorbidities.

Compared to the carbapenem group, the patients treated with the alternative regimen had a shorter hospital stay (median [IQR]: 7 days [5-10] *vs*. 12 days [9-18], p<0.001), (table 1).

In the alternative treatment group, two patients receiving TMP-SMX died due to an ESBL *E. coli* bacteremia (2/25 – 8%), both of which had advanced neoplastic disease (as previously described). This percentage is not higher than the 30 day mortality observed in the carbapenem group (table 1).

### Alternative group

TMP-SMX was the most frequent therapeutic agent selected in these patients. The clinical characteristics and source of bacteremia of patients who received alternative therapy as definitive treatment are shown in table 2, and the complete therapy regimen and length of therapy in the alternative group in table 3. When patients were switched to non-IV antibiotics, they had received a median of 2.5 days (IQR 0-6 days) of intravenous appropriate therapy.

**Table 2.** Clinical characteristics of patients with bloodstream infections caused by extended-spectrum b-lactamaseproducing Enterobacteriaceae who received an alternative therapy as definitive treatment (carbapenem-sparing).

| Patient | Definitive treatment | Sepsis /<br>septic shock | Microorganism | BSI Source | Death | Cure | Relapse |
| --- | --- | --- | --- | --- | --- | --- | --- |
| 1 | TMP-SMX<sup>a</sup> PO<sup>b</sup> / 1 DS<sup>c</sup> tid<sup>d</sup> | No | *E. coli* | Urinary | No | Yes | No |
| 2 | TMP-SMX PO/ 1 DS bid<sup>e</sup> | No | *E. coli* | Urinary | No | Yes | No |
| 3 | TMP-SMX PO/ 1 DS bid | No | *E. coli* | Urinary | No | Yes | No |
| 4 | TMP-SMX PO/ 1 DS bid | Septic shock | *K. pneumoniae amp C* | Catheter | No | Yes | No |
| 5 | TMP-SMX PO/ 1 DS bid | Sepsis | *K. pneumoniae amp C* | Urinary | No | Yes | No |
| 6 | TMP-SMX PO/ 1 DS bid | No | *E. coli* | Biliary | No | Yes | No |
| 7 | TMP-SMX PO/ 1 DS bid | No | *E. coli* | Urinary | No | Yes | No |
| 8 | TMP-SMX PO/ 1 DS bid | No | *E. coli* | Urinary | Yes | No | No |
| 9 | TMP-SMX PO/ 1 DS bid | No | *E. coli* | Urinary | No | Yes | No |
| 10 | TMP-SMX PO/ 1 DS bid | No | *K. pneumoniae amp C* | Biliary | No | Yes | No |
| 11 | TMP-SMX PO/ 1 DS bid | No | *K. pneumoniae amp C* | Urinary | No | Yes | No |
| 12 | TMP-SMX PO/ 1 DS bid | No | *E. coli* | Primary<sup>f</sup> | No | Yes | No |
| 13 | TMP-SMX PO/ 1 DS bid | No | *E. coli* | Urinary | No | Yes | No |
| 14 | TMP-SMX PO/ 1 DS bid | Sepsis | *E. coli* | Urinary | No | Yes | No |
| 15 | TMP-SMX PO/ 1 DS bid | No | *E. coli* | Bowel | No | Yes | No |
| 16 | TMP-SMX PO/ 1 DS bid | No | *E. coli* | Biliary | No | Yes | No |
| 17 | TMP-SMX PO/ 1 DS bid | No | *E. coli* | Urinary | No | Yes | No |
| 18 | TMP-SMX PO/ 1 DS bid | No | *E. coli* | Urinary | No | Yes | No |
| 19 | TMP-SMX PO/ 1 DS bid | Septic shock | *E. coli* | Urinary | No | Yes | No |
| 20 | TMP-SMX PO/ 1 DS bid | No | *K. pneumoniae* | Urinary | No | Yes | No |
| 21 | TMP-SMX PO/ 1 DS bid | Septic shock | *E. coli* | Urinary | No | Yes | No |
| 22 | TMP-SMX PO/ 1 DS bid | No | *E. coli* | Urinary | Yes | No | Yes |
| 23 | TMP-SMX PO/ 1 DS bid | No | *K. pneumoniae* | Primary | No | Yes | No |
| 24 | TMP-SMX PO/ 1 DS bid | No | *E. coli* | Primary | No | Yes | No |
| 25 | TMP-SMX PO/ 1 DS bid | No | *E. coli* | Urinary | No | Yes | No |
| 26 | Fosfomycin PO 500mg qid | No | *K. pneumoniae* | Urinary | No | Yes | No |
| 27 | Fosfomycin PO 500mg qid | No | *E. coli* | Urinary | No | Yes | No |
| 28 | Ciprofloxacin PO 500mg bid | No | *E. coli* | Urinary | No | Yes | No |
| 29 | Ciprofloxacin PO 500mg bid | Septic shock | *E. coli* | Urinary | No | Yes | No |
| 30 | Ciprofloxacin PO 500mg bid | No | *K. pneumoniae Amp C* | Catheter | No | Yes | No |
| 31 | Ciprofloxacin PO 500mg bid | No | *K. pneumoniae* | Catheter | No | Yes | No |
| 32 | Ciprofloxacin PO 500mg bid | No | *E. coli* | Urinary | No | Yes | No |
| 33 | Ciprofloxacin PO 750mg bid | No | *E. coli* | Catheter | No | Yes | No |
| 34 | Ciprofloxacin PO 500mg bid | No | *E. coli* | Urinary | No | Yes | No |
| 35 | Ciprofloxacin PO 500mg bid | No | *E. coli* | Abdominal | No | Yes | No |
| 36 | Ciprofloxacin PO 500mg bid | No | *E. coli* | Urinary | No | Yes | No |
| 37 | Amoxicillin/clavulanate ER<sup>g</sup><br>PO 1000/62.5 mg qid | No | *E. coli* | Urinary | No | Yes | No |
| 38 | Gentamicin IM<sup>h</sup> 240 mg qd | No | *E. coli* | Urinary | No | Yes | No |
| 39 | Gentamicin IM 240 mg qd | Sepsis | *E. coli* | Urinary | No | Yes | No |
| 40 | Gentamicin 240 mg qd | No | *E. coli* | Urinary | No | Yes | No |
| 41 | Amikacin IM 1 g qd | No | *E. coli* | Urinary | No | Yes | No |
| 42 | Gentamicin 240 mg qd | No | *E. coli* | Urinary | No | Yes | No |

<sup>a</sup>TMP-SMX: Trimethoprim-Sulfamethoxazole. <sup>b</sup>PO: Oral administration. <sup>c</sup>DS: double strength. <sup>d</sup>tid: three times a day. <sup>e</sup>bid: twice a day <sup>f</sup>Primary: No source identified. <sup>g</sup>ER: extended release. <sup>h</sup>IM: intramuscular administration. <sup>i</sup>qd: once daily.

**Table 3.**
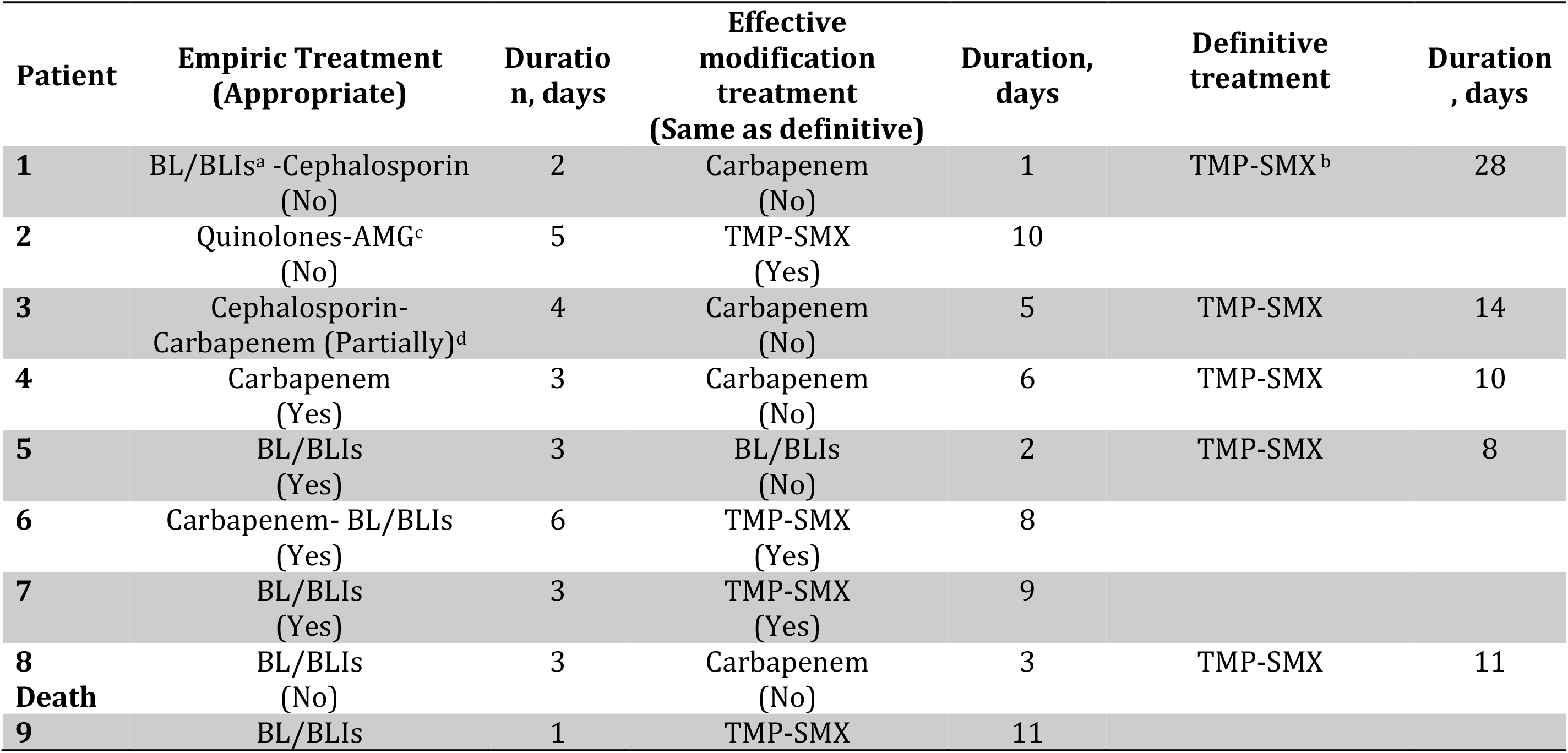
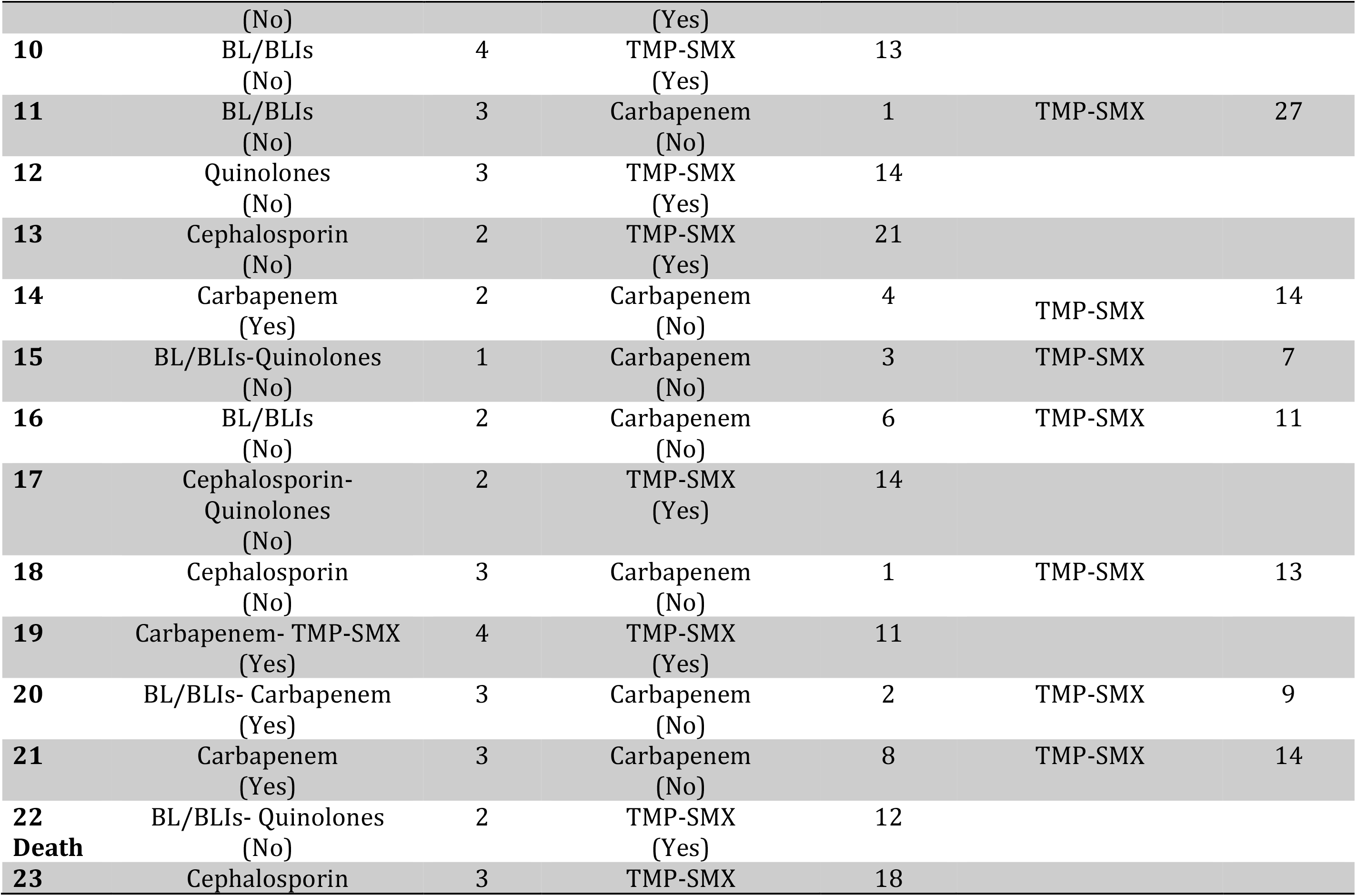
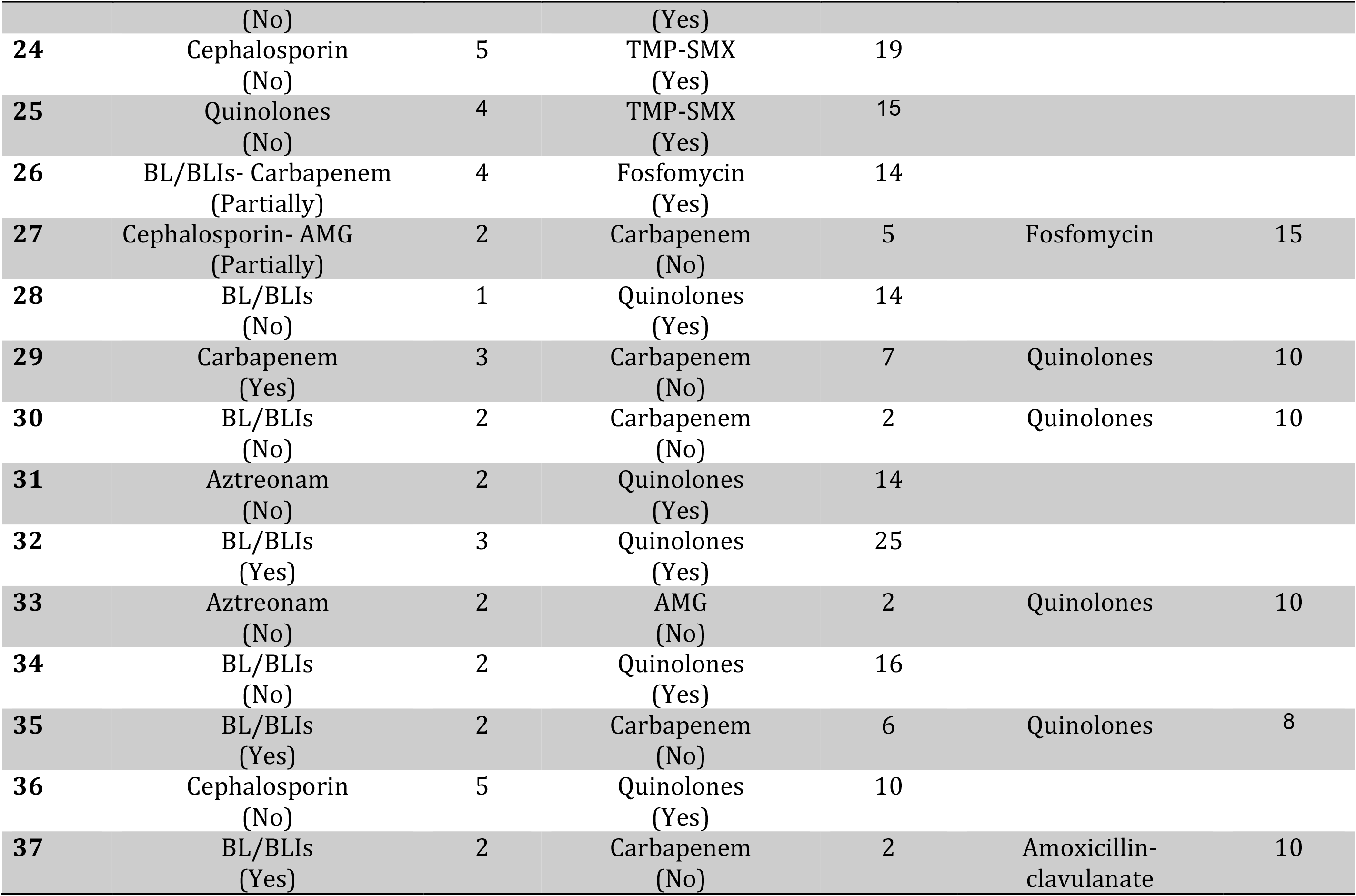
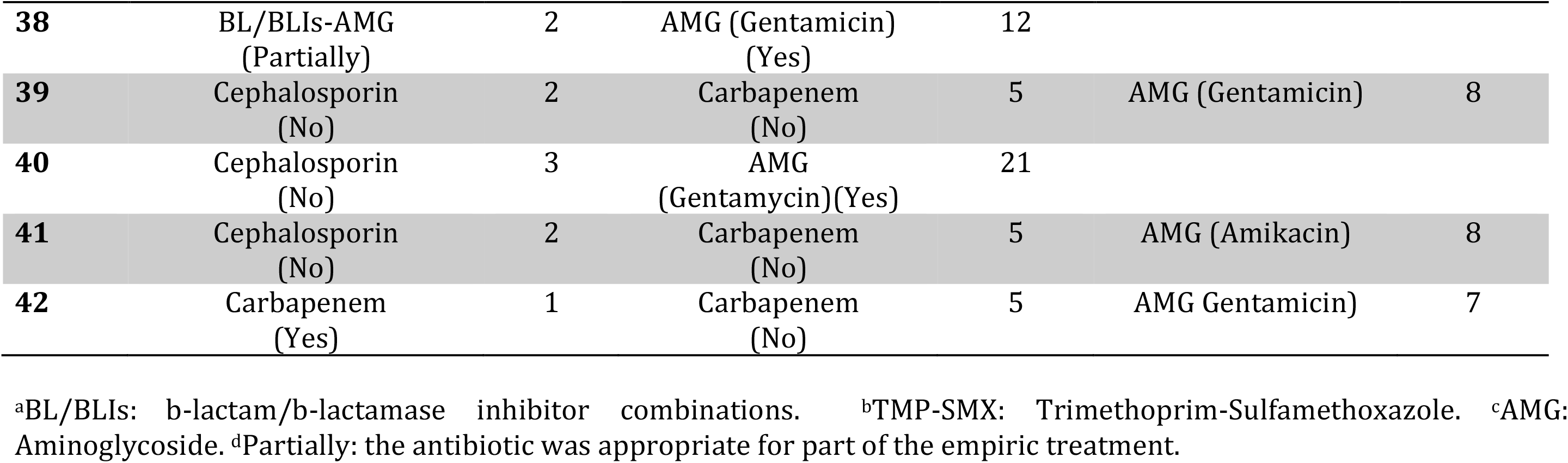
Complete therapy regimen and duration of treatment of patients with bloodstream infections caused by extended-spectrum b-lactamase-producing Enterobacteriaceae who received an alternative therapy as definitive treatment (carbapenem-sparing).

### Multivariate and propensity score analysis

In the univariate analysis, nosocomial acquisition (OR 4.08; 95% CI 1.10-15.11; p=.035), chronic kidney disease (OR 6.22; 95% CI 1.53-25.27; p=.01) and the microorganism *(K. pneumoniae* compared with *E. coli)* (OR 3.85; 95% CI 1.05-14.20; p=.04), were independent predictors of clinical failure. The use of an alternative non-IV treatment was not related to mortality (OR 0.27; 95% CI 0.05-1.61; p=.15) (table 4). After controlling for confounding with the propensity score, the adjusted OR of carbapenem treatment was 4.95; 95% CI (0.94-26.01, p=.059), (table 5).

**Table 4.** Univariate and multivariate analysis. Relation between variables and failure.

| • UNIVARIATE ANALYSIS |  | OR | (95%CI) | p-value |
| --- | --- | --- | --- | --- |
| Age |  | 0.99 | ( 0.94; 1.03) | 0.57 |
| Sex (male) |  | 1.79 | ( 0.44; 7.14) | 0.42 |
| Diabetes Mellitus |  | 0.43 | ( 0.05; 3.59) | 0.43 |
| Chronic kidney disease |  | 6.22 | ( 1.53; 25.27) | **0.011** |
| Immunosuppression |  | 2.00 | ( 0.37; 10.73) | 0.42 |
| Cancer |  | 1.23 | ( 0.30; 5.07) | 0.77 |
| Charlson Index |  | 1.08 | ( 0.86; 1.35) | 0.49 |
| Sepsis / Septic Shock |  | 0.33 | ( 0.04; 2.72) | 0.30 |
| Pitt score |  | 0.84 | ( 0.54; 1.29) | 0.42 |
| Nosocomial acquisition |  | 4.08 | ( 1.10; 15.11) | 0.03 |
| Health-care associated |  | 1.94 | ( 0.55; 6.92) | 0.30 |
| ESBL microorganism | *K. pneumoniae* | 3.85 | ( 1.05;14.20) | **0.042** |
| Bloodstream infection source <sup>1</sup> | Urinary tract | 1 |  | 0.52 |
|  | Catheter related | 1.000 | ( 1.00; 1.00) |  |
|  | Biliary tract | 2.42 | ( 0.53; 11.04) |  |
|  | Other | 1.38 | ( 0.25; 7.59) |  |
| Appropriate empirical therapy | Yes | 0.70 | ( 0.18; 2.66) | 0.84 |
|  | Partially<sup>b</sup> | 1.19 | ( 0.12; 11.71) |  |
| Delay of appropriate therapy <sup>a</sup> |  | 1.07 | ( 0.73; 1.57) | 0.74 |
| Empirical treatment duration in days |  | 1.004 | ( 0.58; 1.72) | 0.99 |
| Definitive treatment duration in days |  | 0.98 | ( 0.89; 1.09) | 0.75 |
| Alternative non-IV treatment |  | 0.28 | ( 0.06; 1.35) | 0.11 |
| Hospital stay after BSI <sup>b</sup> |  | 1.01 | ( 0.99; 1.04) | 0.34 |
| • MULTIVARIATE ANALYSIS |  | OR | (95%CI) | p-value |
| Chronic kidney disease |  | 11.19 | (1.84; 67.94) | 0.009 |
| ESBL microorganism ( *K. pneumoniae*) |  | 7.86 | (1.16-53.41) | 0.035 |
| Alternative non-IV treatment |  | 0.27 | (0.05-1.61) | 0.15 |

<sup>a</sup> Delay of appropriate therapy: days without active treatment in terms of antibiotic susceptibility.
<sup>b</sup> BSI: bloodstream infection

**Table 5.**
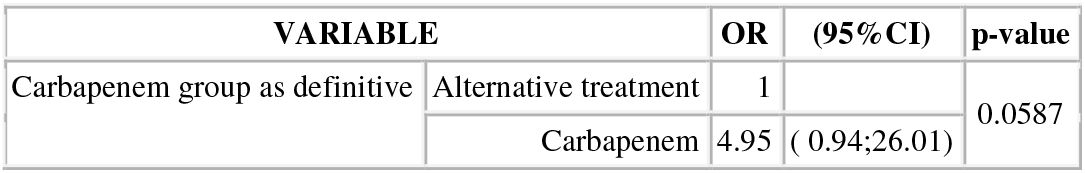
Propensity score analysis.

## DISCUSSION

In this study we have observed that the use of alternative non-IV carbapenem-sparing antibiotics for definitive treatment of ESBL or Amp-C-positive Enterobacteriaceae bloodstream infections was not related to greater mortality. In fact we did not find differences in either the primary outcome, 30-day mortality (6 [10%] *vs*. 2 [5%], p=0.46) nor the secondary outcome, clinical failure (9 [15%] vs. 2 [5%], p=0.12) for carbapenem group *vs*. non-IV carbapenem-sparing antibiotics. Moreover, the use of alternative treatment was associated with a shorter hospital stay.

Some reports have evaluated the efficacy of IV carbapenem-sparing antibiotics in this setting, including cephamycins, BL/BLIs or fluoroquinolones, and have presented both positive (5, 9, 12, 13, 23) and negative outcomes (24, 25). In a metanalysis (3), the use of empirical quinolones (oral or intravenous) for ESBL Enterobacteriaceae bacteremia was associated with a higher mortality than carbapenems, but mortality was similar when quinolones were used as definitive therapy. However, even in the carbapenem-sparing setting, some studies have shown prior exposure to fluoroquinolones or β-lactam to be independent risk factors for ESBL or carbapenem-resistant enterobacteriae infections (26, 27). Therefore, these antibiotics would probably not be the best options for the treatment of these infections.

To our knowledge, there is no published experience with other oral alternatives such as TMP-SMX or fosfomycin for the treatment of ESBL Enterobacteriaceae bacteremia (28). Published experience with TMP-SMX for the treatment of ESBL or Amp-C infections in patients without bacteremia is also scarce. Park *et al*. showed that non-carbapenem antibiotics (which include 5 patients treated with TMP-SMX) had a similar efficacy to carbapenems among a case series of pyelonephritis, however the outcome of patients treated with TMP-SMX was not specified (29). TMP-SMX was the most frequent option chosen as non-IV, carbapenem-sparing, definitive treatment in our study (mainly for urinary and biliary sources); no complications were found related to this use. Our experience with TMP-SMX, after confirming antibiotic susceptibility (38% of the strains of ESBL infections in our setting), is promising. This option may prevent the emergence of resistance, it allows for the administration of an oral regimen, and it could shorten the hospital stay.

Studies of the efficacy of non-IV carbapenem-sparing agents for infections caused by ESBL/Amp-C-producing Enterobacteriaceae have focused mainly on urinary tract infections (30, 31), have not assessed cases of bacteremia, and have addressed mainly patients with *ESBL-E.coli* infections. Since most of the available drugs (TMP-SMX, quinolones, fosfomycin) have high urinary levels, further studies should determine if this alternative non-IV carbapenem-sparing agents are also useful for the treatment of other bacteremic foci (abdominal) and for *ESBL-Klebsiella* infections.

Data on non-IV carbapenem-sparing treatments could help reduce carbapenem use, which is crucial in order to contain the spread of carbapenem resistance (32), to reduce its impact on global hospital ecology (33), and to shorten hospital stays. As demonstrated, hospital stays in the alternative treatment group were significantly shorter in our study population without a negative impact in terms of relapse or early re-admission. The benefits associated with non-prolonged hospitalization in terms of cost-effectiveness and comorbidity have been already demonstrated (34).

This study has certain limitations. The retrospective design could not exclude the possibility that patients with more severe infections were preferably treated with carbapenems without subsequent treatment with a non-carbapenem. All cases were included in this retrospective study. Further, the sample size may be too small to achieve adequate statistical power and selection by indication may bias the results. However we tried to balance this limitation by adjusting our results using a propensity score analysis, and did not observe changes in estimation effects. This study may not account for all the variables that may have influenced the decision to use carbapenems and thus might influence the OR; similarly, the goodness of model fit for calculating propensity score weights might be underpowered (AUC=.77). We also could not characterize the ESBL genes or investigate the MIC distribution for all study isolates. Finally, the study included mostly bloodstream infections due to *E. coli* which means that these results cannot be extrapolated to *K. pneumoniae* and mixed ESBL/amp-C-positive Enterobacteriaceae or other species of the Enterobacteriaceae family.

In spite of these limitations, the possibility that non-IV antibiotics could be used for the definitive treatment of (ESBL)-Amp-C-positive Enterobacteriaceae bloodstream infections is promising. Our data supports the use of TMP-SMX as a carbapenem-sparing alternative therapy which could reduce carbapenem use and shorten hospital stays. Larger prospective interventional studies are now required to definitively assess the efficacy of oral carbapenem-sparing antibiotics for the treatment of ESBL-Amp-C-positive Enterobacteriaceae bacteremia.

## ACKNOWLEDGMENTS

We thank Jay A Fishman for his critical review, Santi Pérez Hoyos and Alex Sànchez for statistical study and Michael Maudsley for English language support and editorial assistance. CP, NF-H and JR-B are supported by Plan Nacional de I+D+i 2013-2016 and Instituto de Salud Carlos III, Subdirección General de Redes y Centros de Investigación Cooperativa, Ministerio de Economía, Industria y Competitividad, Spanish Network for Research in Infectious Diseases (REIPI RD16/0016/0001 and 0003), which is co-financed by European Development Regional Fund “ A way to achieve Europe “, Operative Programme Intelligent Growth 2014 – 2020.

## TRANSPATENCY DECLARATION

The authors have no conflicts of interest to declare.

